## Supplementary Figures for "Minimal background noise enhances neural speech tracking: Evidence of stochastic resonance"

Björn Herrmann*

Rotman Research Institute,
Baycrest Academy for Research and Education, M6A 2E1, North York, ON, Canada

Department of Psychology,
University of Toronto, M5S 1A1, Toronto, ON, Canada

*Correspondence concerning this article should be addressed to Björn Herrmann, Rotman Research Institute, Baycrest, 3560 Bathurst St, North York, ON, M6A 2E1, Canada.

**Keywords:** Electroencephalography, speech encoding, story listening, temporal response function, background noise, attention

### Supplementary Results

#### P1-N1 amplitudes using the amplitude envelope of speech

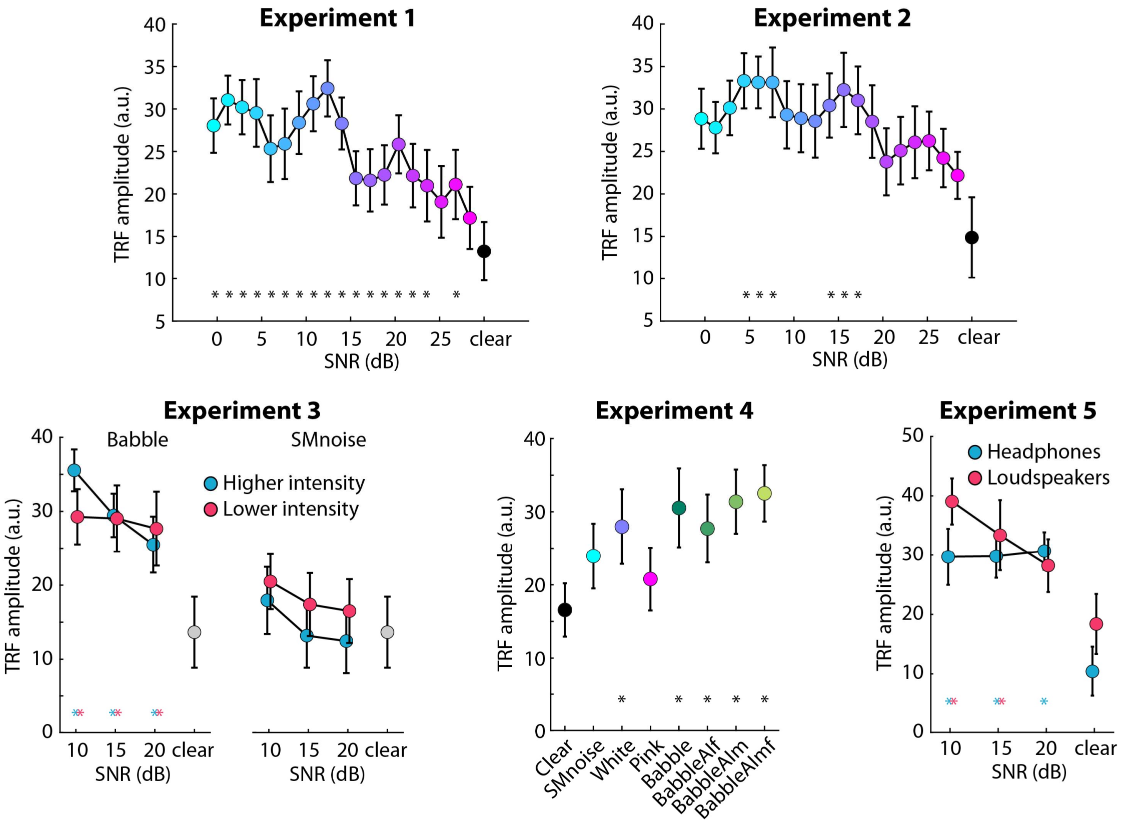


**Figure 1-figure supplement 1: P1-N1 amplitude from TRF analyses using the amplitude envelope of speech.** An asterisk close to the x-axis indicates a significant difference relative to the clear condition (FDR-thresholded). The absence of an asterisk indicates that there was no significant difference. Error bars reflect the standard error of the mean. For additional details, see the respective figure captions in the main article.

#### No relationship between visual-task performance and neural speech tracking in Experiment 2

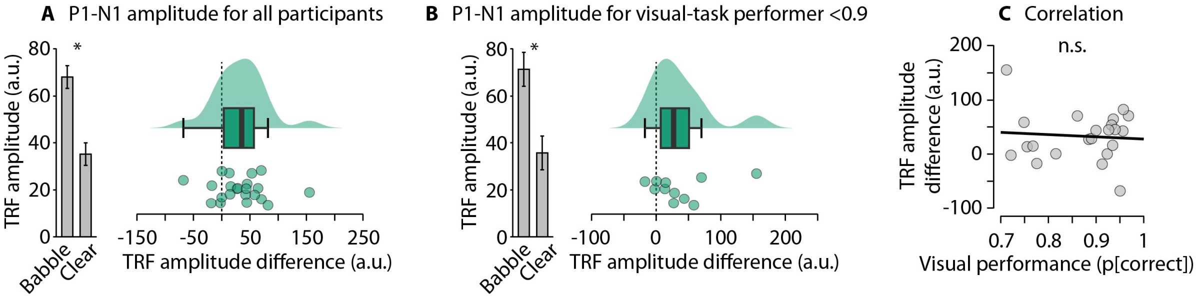


**Figure 2-figure supplement 1: Relationship between visual performance and the noise-related enhancement of the P1-N1 amplitude in Experiment 2. A:** Shows the P1-N1 amplitude for speech in babble (averaged across all SNRs above 15 dB, for which speech was highly intelligible) and clear speech. **B:** Same as in Panel A for individuals performing below 0.9 in the visual task(to controls for potential influences of high performers, who could have attended the speech). **C:** Correlation between visual-task performance and the difference in the P1-N1 amplitude between speech in babble and clear speech. The relationship was not significant. *p < 0.05, n.s. – not significant.

#### P1-N1 amplitudes using cross-correlation instead of temporal response function

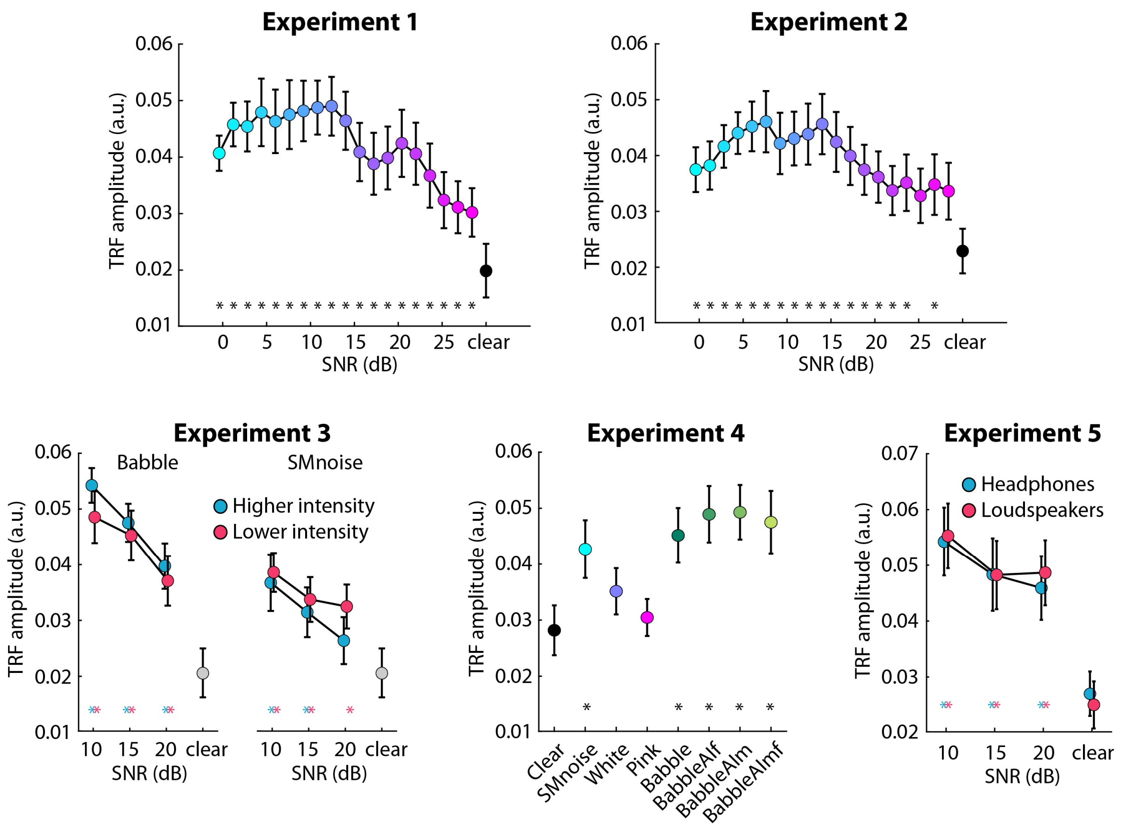


**Figure 1-figure supplement 2: P1-N1 amplitude from cross-correlations analyses.** An asterisk close to the x-axis indicates a significant difference relative to the clear condition (FDR-thresholded). The absence of an asterisk indicates that there was no significant difference. Error bars reflect the standard error of the mean. For additional details, see the respective figure captions in the main article.
